# Reverse-engineering amyloid strains with generative protein design

**DOI:** 10.64898/2026.05.08.723915

**Authors:** Laxmikant Gadhe, Katerina Konstantoulea, Aneesh Mazumder, Jie Chen, Lukasz A. Joachimiak, Nikolaos N. Louros

## Abstract

Amyloid fibrils are intrinsically polymorphic protein assemblies that form distinct structural strains linked to diverse biological and pathological outcomes. Yet, the principles governing how sequence encodes diverse fibril architectures, and the extent to which a given fold constrains underlying amino-acid sequence compatibility, remain poorly understood. Here, we apply generative protein design to directly interrogate the sequence–structure relationship of defined fibril architectures, using α-synuclein (αS), a protein known to form highly polymorphic amyloid fibrils, as a model system. Sampling sequence space under structural constraints reveals a continuous compatibility manifold in which diverse sequences encode a common amyloid architecture. *De novo* designed sequences assemble into fibrils, often with enhanced aggregation efficiency relative to αS. A subset exhibits strain-like behaviour, including similar morphologies, efficient cross-templating, and induction of αS cellular propagation, thereby functionally validating structural compatibility with the native fibril fold. Energetic analysis shows that stability is achieved through distinct but compensatory interactions, supporting a non-unique mapping between sequence and structure. Together, our results define a continuous and constrained compatibility landscape underlying amyloid strains, providing a framework for understanding the determinants of polymorphism and establishing generative protein design as a strategy to access this space, interrogate amyloid sequence–structure relationships, and engineer fibrillar protein assemblies and functional biomaterials.

## Introduction

Amyloid fibrils are highly ordered protein assemblies formed by a wide range of proteins and are a defining feature of numerous neurodegenerative and systemic diseases^5^. These assemblies share a common cross-β architecture^6^, yet can adopt multiple structurally distinct polymorphs, often referred to as amyloid strains, which are associated with differences in disease phenotype, progression, and tissue specificity^3,7,8^. Understanding how protein sequence encodes these diverse structural states remains a central challenge in amyloid biology. Amyloid fibrils formed by α-synuclein (αS) exhibit remarkable structural polymorphism^9,10^ and constitute defining pathological hallmarks of several neurodegenerative disorders collectively known as synucleinopathies^11,12^, including Parkinson’s disease (PD)^13^, dementia with Lewy bodies^14,15^, and multiple system atrophy (MSA)^16^. High-resolution cryo-electron microscopy (cryo-EM) studies have revealed multiple αS fibril polymorphs derived from both *in vitro* assemblies^17^ and patient brain material^18–20^, demonstrating that distinct folds can arise from the same primary sequence across different disease contexts, yet are conserved among patients with the same pathology. These structural differences are thought to underlie the pathological diversity observed across synucleinopathies, highlighting the importance of understanding the relationship between sequence, structure, and strain behavior^21,22^.

Despite these advances, a fundamental question remains unresolved: to what extent does a given amyloid fold constrain the underlying amino-acid sequence? In globular proteins, many structural scaffolds can accommodate a wide range of sequences provided that key stabilizing interactions are preserved. Whether similar principles apply to amyloid fibrils is less clear. Thermodynamic^2,4^ and geometric^23–25^ analyses of amyloid structures have shown that steric zipper interactions between short aggregation-prone regions stabilize the fibril core and impose constraints on side-chain packing, while also suggesting a limited but non-zero tolerance for sequence variation within defined structural contexts^1,9,25^. Consistent with this, naturally occurring mutations and posttranslational modifications can alter aggregation behaviour and fibril structure without abolishing fibril formation^18,26,27^. Determining the sequence tolerance of a defined amyloid fold is therefore critical for understanding the molecular determinants of fibril assembly, the emergence of polymorphic diversity, and the potential to engineer amyloid scaffolds as functional materials.

Recent advances in computational protein design provide an opportunity to directly interrogate sequence–structure compatibility in amyloid architectures. Deep learning– based generative design approaches have matured into powerful tools capable of inferring molecular grammars that enable efficient exploration of sequence space compatible with predefined structural backbones^28,29^, designing novel protein folds^30–34^, and engineering customised proteins with atomic-level accuracy^35–39^. While these methods have been widely applied to globular proteins, their extension to amyloid fibrils remains unexplored. Amyloid assemblies differ fundamentally from globular proteins: their interfaces are repetitive and modular, stability is distributed across stacked monomers rather than compact domains^3,4^, and pathological strains are not constrained by evolutionary selection in the same way as folded proteins^40^. Nevertheless, the availability of high-resolution fibril structures provides a unique opportunity to use these architectures as templates to systematically probe the sequence constraints required to maintain a given amyloid fold.

Here, we extend structure-guided generative protein design to amyloid fibrils to directly probe the sequence–structure relationship of αS assemblies and to reverse-engineer *de novo* amyloid strains compatible with defined fibril architectures (**Fig. 1**). Using previously determined αS fibril structures as templates, we systematically explored the sequence space of their fibril core while preserving solvent-exposed residues of the wild-type (WT) protein. Designed sequences were evaluated computationally using energetic modelling and stability analysis, and selected candidates were recombinantly produced for experimental characterization. We show that *de novo* designed sequences readily assemble into amyloid fibrils, with a subset reproducing key features of the template strain, including spectral signatures, cross-templating of αS, and propagation in cellular biosensor assays. Together, these findings demonstrate that amyloid folds define a continuous but constrained sequence compatibility regime and establish generative protein design as a framework for interrogating and reprogramming amyloid strain behaviour from existing structural templates.

**Figure 1.**
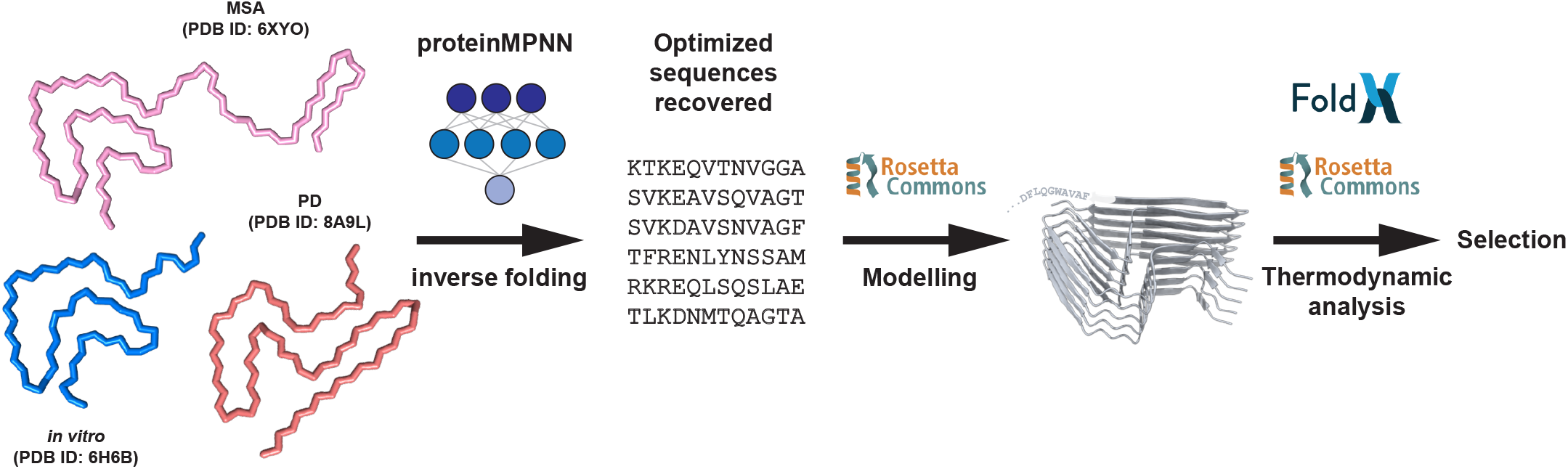
Reverse-engineering strategy of *de novo* designs compatible with amyloid folds. Schematic of the computational design workflow. Experimentally determined fibril structures are used as a template to explore sequence compatibility of the fibril core using deep-learning-guided inverse folding. The resulting sequences are then threaded back to the template structure, and the derived structural models are employed for thermodynamic evaluation and ranking.

## Results

### A conserved sequence–structure compatibility landscape across fibril templates

Amyloid fibrils formed by αS exhibit substantial structural polymorphism, raising the question of whether sequence compatibility is specific to individual fibril architectures or reflects a more general property of amyloid folds. To address this, we applied structure-guided sequence design to three distinct αS fibril structures, including an MSA-derived model (PDB ID: 6XYO)^20^, an *in vitro* derived fibril structure (PDB ID: 6H6B)^17^, and the structure of PD patient-derived fibrils (PDB ID: 8A9L)^19^ (**Fig. 2A**). For each template, inverse folding was performed under symmetry constraints to simultaneously mutate stacked fibril core positions, with solvent-exposed residues and the less conserved NAC-residue containing segment of the fibril core held fixed (**Fig. 2A**, segment **B**). In contrast, residues within the remainder of the buried fibril core were allowed to vary (**Fig. 2A**, segment A). Inverse folding used to sample sequences across a range of temperatures enabled controlled exploration of sequence space under progressively relaxed design constraints. Alignment of designed sequences from the unconstrained regime revealed that patient-associated αS fibril structures impose stronger constraints than the *in vitro*-derived structure. In particular, the disease-associated templates retained higher conservation across more positions, suggesting a narrower sequence–structure compatibility space due to stricter packing constraints. Critical glycine residues were recovered across polymorphs, consistent with strict geometric requirements at these sites (**Fig. 2B**). When design was restricted to the A segment of the fibril core, the three templates showed broadly similar levels of conservation across the designed positions. This suggests that, under matched structural constraints, the accessible sequence space of this region is comparable across αS polymorphs. However, the residue identities favoured by each template differed. Patient-derived structures were enriched in hydrophobic substitutions, whereas the *in vitro* template tolerated more polar and charged residues within the fibril core (**Fig. 2B**). This indicates that similar levels of sequence variability can arise from distinct physicochemical solutions depending on the local packing environment of each fibril architecture.

**Figure 2.**
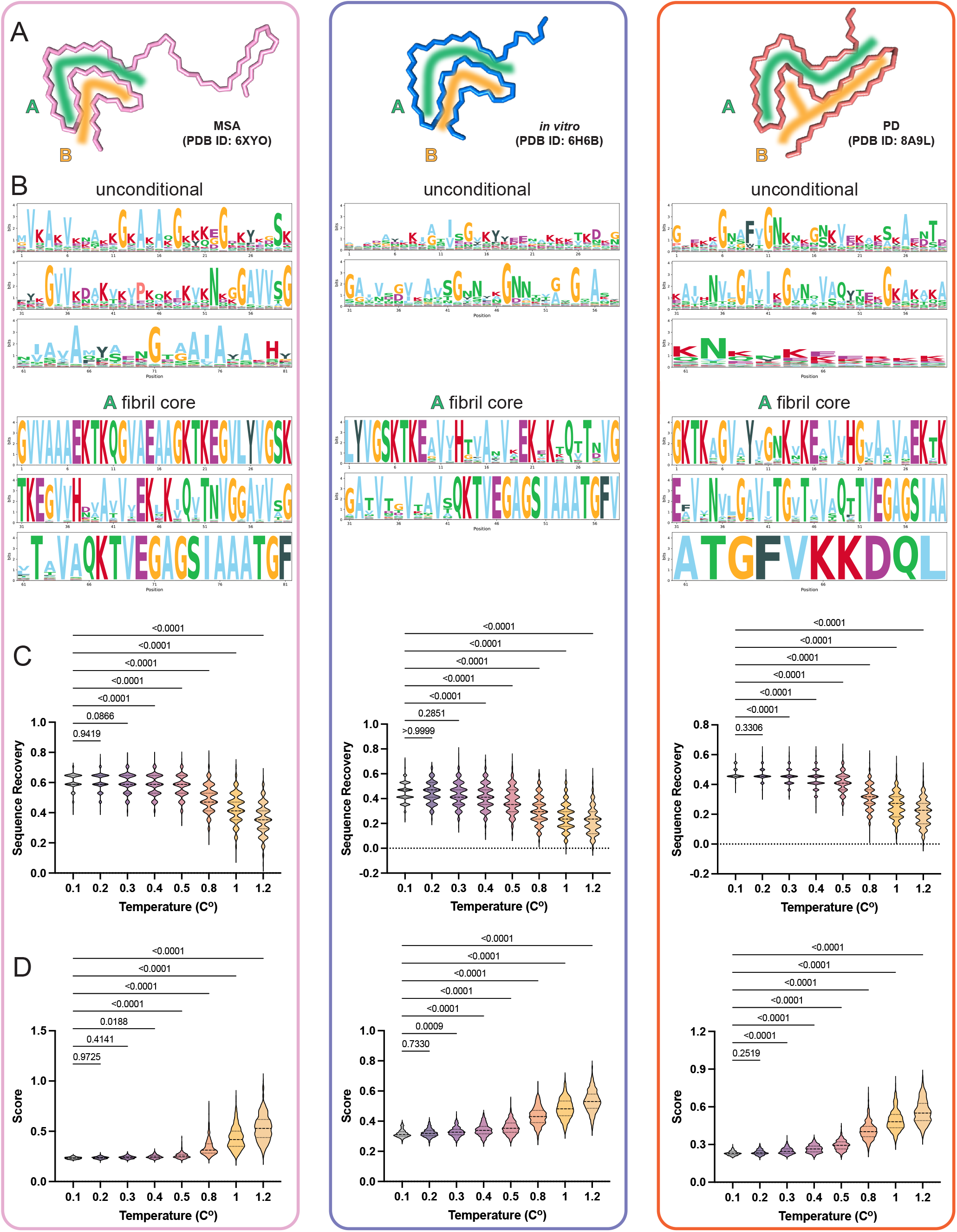
Structure-guided generative design across αS fibril templates. (**A**) Schematic of the design strategy. Cryo-EM structures of αS fibrils representing an MSA-derived model (PDB ID: 6XYO), an *in vitro*–derived fibril (PDB ID: 6H6B), and a patient-derived fibril (PDB ID: 8A9L) were used as templates for inverse folding sequence design (n=500 sequences per temperature and per template). (**B**) Sequence alignment logos derived from unconstrained and A-segment-restricted design regimes. (**C**) Sequence recovery as a function of sampling temperature for each structural template. (**D**) Distribution of design scores across sampling temperatures. Statistics: One-way ANOVA with Dunnett correction for multiple comparisons.

Analysis of sequence recovery and design scores across sampling temperatures revealed a consistent trade-off across all templates (**Fig. 2C-D**). Sampling temperature further modulated this compatibility landscape, as sequence recovery decreases sharply beyond T ≈ 0.4–0.5, while design scores increase monotonically, reflecting progressive exploration of sequence space. Notably, this transition regime is conserved across all three templates, indicating that the balance between sequence conservation and structural compatibility is a general property of αS amyloid sequence design rather than a feature of a specific fibril architecture. Together, these analyses indicate that αS folds impose both shared and template-specific sequence constraints: disease-associated structures appear more restrictive under unconstrained design, whereas targeted design of the conserved A segment reveals comparable overall tolerance but distinct physicochemical preferences across polymorphs.

### Temperature-dependent drift along a continuous sequence compatibility manifold

To characterize how designed sequences populate this generated landscape, we analysed their aggregation propensity and physicochemical properties across sampling temperatures (**Fig. 3**). Aggregation propensity profiling revealed a strong anti-correlation between TANGO and WALTZ scores across temperatures (**Fig. 3A**). Increasing sampling temperature leads to a decrease in TANGO-derived aggregation propensity, which captures broadly hydrophobic, β-sheet-driven aggregation^41^, and a concomitant increase in WALTZ scores, which reflect sequence-specific amyloid-forming motifs^42^. This is further highlighted by tracking overall change in secondary structure propensities, with a gradual shift from β-to α-propensity (**Fig. 3B**). This suggests that sampling temperature does not simply reduce aggregation propensity. Instead, under structural constraints, sampling explores sequence space that deviates from generic hydrophobic-driven aggregation propensities toward sequences structurally compatible with the cross-β amyloid architecture, consistent with previous studies showing that structure-based approaches can improve prediction of amyloid motifs^43^. Consistent with this, analysis of predicted aggregation-prone windows (**Fig. 3C-D**) and hydropathy profiles (**Fig. 3E**) revealed a temperature-dependent design drift. Dimensionality reduction of the engineered sequence space using physicochemical residue descriptors further showed that these changes occur gradually rather than through discrete transitions, as projection into the latent space revealed that the designed sequences occupy a continuous manifold rather than forming discrete clusters (**Fig. 3F**). Interestingly, although higher temperatures generate sequences with decreased similarity to the WT, as indicated by their reduced average BLOSUM62 scores (**Fig. 3G**), the overall physicochemical composition of the engineered sequences remains consistent across temperatures. The WT sequence is positioned centrally within this latent space (**Fig. 3G**, black), suggesting that sequence divergence derived at higher temperatures does not significantly alter the physicochemical composition and potential compatibility with the original template scaffold. However, the distribution of sequences shows a clear temperature-dependent directionality within the manifold.

**Figure 3.**
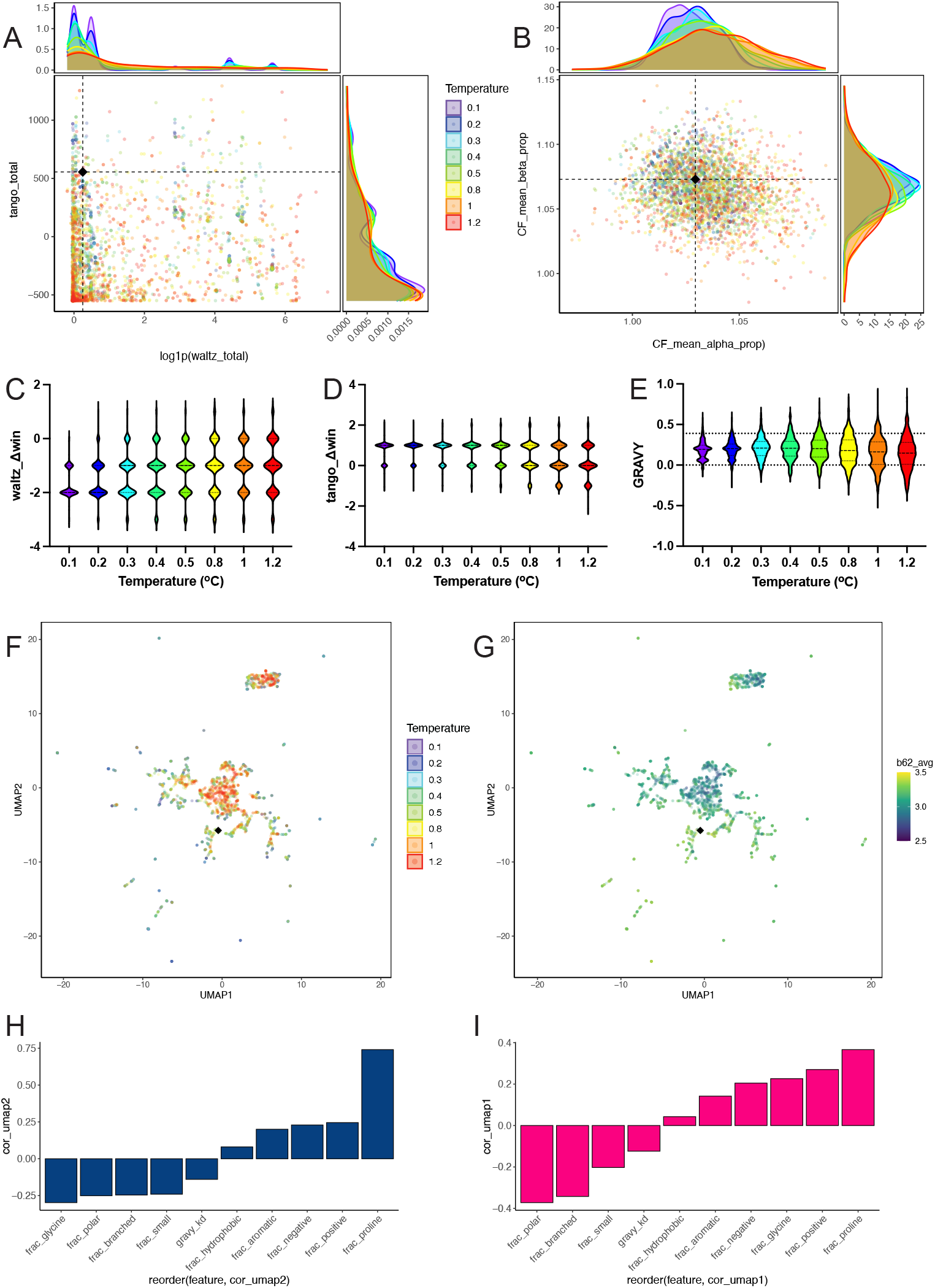
Exploration of the sequence-structure compatibility design landscape. (**A**) Correlation analysis between TANGO and WALTZ aggregation propensity scores across designed sequences, showing anti-correlation across sampling temperatures. (**B**) Predicted secondary structure propensities as a function of sampling temperature, indicating a gradual shift from β-to α-propensity. (**C–D**) Distribution of aggregation-prone region windows predicted by TANGO and WALTZ, and (**E**) hydropathy scores of designed sequences across sampling temperatures. (**F**) UMAP projection of designed sequences based on physicochemical descriptors, showing a continuous sequence manifold with temperature-dependent directionality. (**G**) UMAP projection indicating sequence similarity to αS (BLOSUM62 score). (**H–I**) Correlation of physicochemical features with UMAP axes.

Analysis of feature contributions to this embedding suggests that this variation along the principal axes is driven by coordinated changes in residue composition, rather than by a single dominant feature (**Fig. 3H-I**), with higher sampling temperatures exploring increased charge and aromatic content, and lower temperatures favouring polar and branched amino acid compositions.

Together, these results define a low-dimensional sequence compatibility manifold governed by the structural constraints of the template amyloid fold. This indicates that substantial sequence divergence can be achieved without exiting the physicochemical regime compatible with the αS fibril architecture.

### Energetic evaluation of *de novo* engineered amyloid cores

To evaluate the structural compatibility of designed sequences with the target fibril architecture, we first assessed global energetics using Rosetta-based modelling. Designed sequences were threaded onto the fibril template structure and subjected to structural relaxation, and their total energies were used as a proxy for compatibility with the fold. Analysis of Rosetta scores across sampling temperatures revealed a progressive broadening of the energy distribution at higher temperatures (**Fig. 4A**). While the average energy increases with temperature, reflecting the inclusion of less structurally compatible sequences, this regime also gives rise to a subset of designs with particularly favourable energies. This suggests that higher-temperature sampling not only introduces non-compatible sequences but also enables exploration of more distant areas of sequence space, from which alternative combinations of residues can potentially stabilize the fibril core.

**Figure 4.**
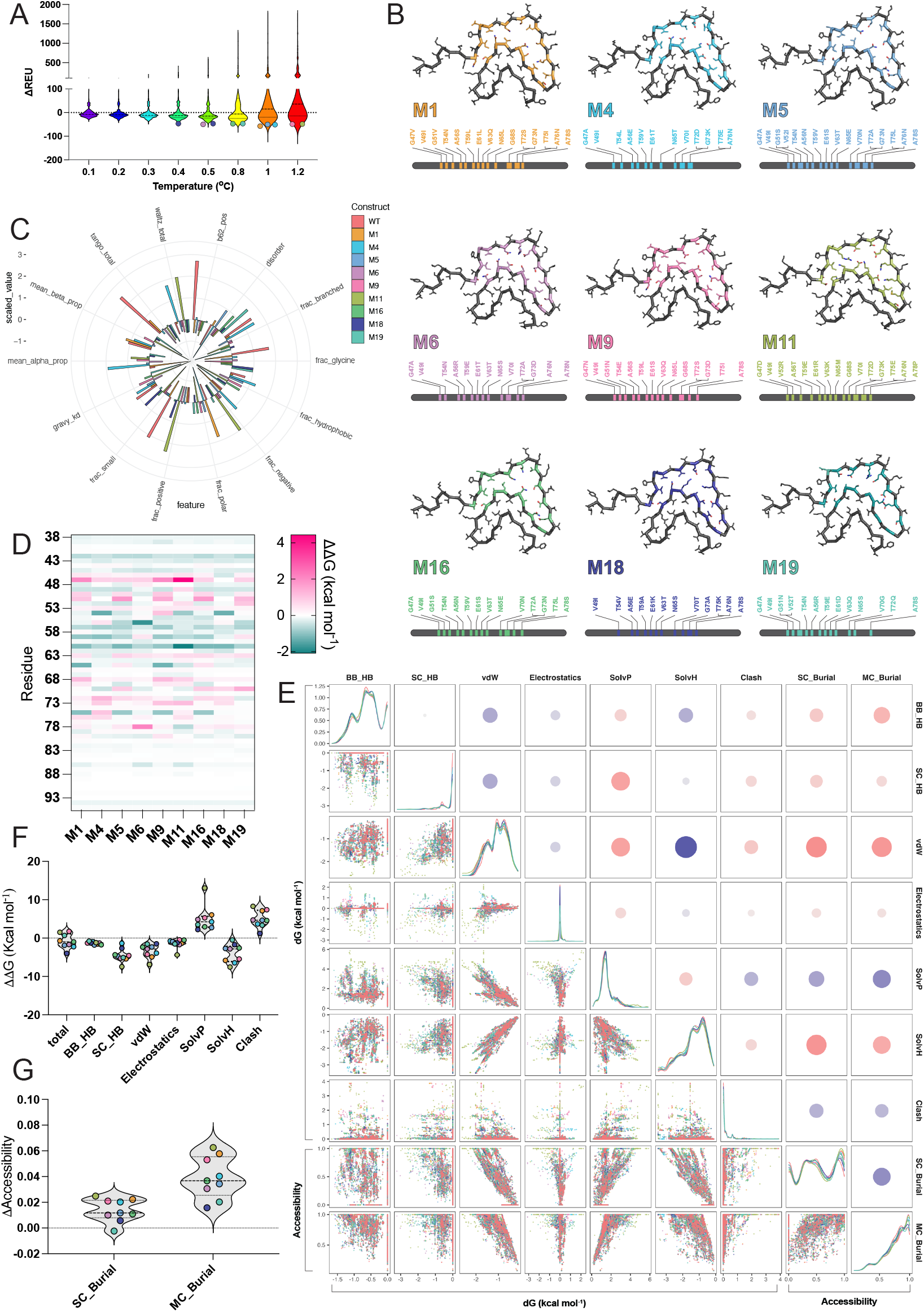
Energetic analysis reveals multiple sequence solutions compatible with a single fibril fold. (**A**) Distribution of Rosetta energies for designed sequences across sampling temperatures, showing increased variance at higher temperatures. Sequences selected for further analysis (n=9) are shown as coloured points, matching colours shown in (B). (**B**) Selected representative designs from higher-temperature regimes used for detailed energetic analysis. (**C**) Circular histogram of scaled physicochemical descriptors for selected αS designs, matching colours shown in (B), relative to the WT sequence. (**D**) Per-residue ΔΔG profiles relative to the αS template, calculated using established profiling methods^1–4^, highlight distributed stabilizing and destabilizing contributions across the fibril core. (**E**) Global stability comparison of *de novo* designs relative to the αS template structure. (**F**) Correlation analysis of energy components, showing coupling between van der Waals interactions, solvation, and electrostatics. (**G**) Side-chain and main-chain accessibility across designs, indicating modest improvement in packing.

To further investigate the energetic basis of these observations, we selected representative low-scoring sequences from higher sampling temperatures (T ≥ 0.4), corresponding to the regime in which sequence divergence becomes more pronounced (**Fig. 4B**). Scaling and inspection of their physicochemical properties revealed that these sequences do not arise from optimization of a single feature but instead occupy a constrained multi-dimensional region of sequence space, with most features centred near the baseline and exhibiting directional perturbations (**Fig. 4C**). Homology to the WT is low across the designed space, where fractions of branched, glycine and short aliphatic chains are reduced, significantly impacting overall hydrophobicity, β-propensity and hydrophobicity-based aggregation propensity. As expected, designs derived from higher sampling temperatures show stronger deviations along individual axes. This pattern indicates that structural compatibility could be governed by coordinated constraints across multiple sequence features rather than by a single dominant determinant, consistent with the distributed nature of stabilizing interactions within amyloid fibrils, where side-chain packing, hydrogen bonding, and solvation effects collectively define the energetic landscape. As a result, multiple distinct sequences can satisfy the geometric and energetic requirements of the fold through different combinations of local interactions, provided that global physicochemical balance is maintained. These observations define a “compatibility regime” within which sequence variation is tolerated and from which experimentally viable designs can be selected.

These sequences were subjected to detailed residue-level energetic analysis using FoldX, which evaluates the energetic impact of sequence variation by introducing mutations onto a fixed backbone and optimizing sidechain rotamers. This approach enables direct assessment of sequence compatibility within the geometric constraints of the original fibril architecture, providing a complementary perspective on the Rosetta-based energetic determinants of stability. Profiling per residue contributions to fibril stability, compared to the WT αS sequence, revealed a relatively homogeneous distribution across the fibril core (**Fig. 4D**), with residues within the redesigned segment showing improved stability, often accompanied by compensatory destabilization of flanking positions to accommodate changes in the fibril core. Despite substantial sequence divergence, most designs exhibit improved global stability relative to the WT αS structure, with no evidence of large-scale destabilization (**Fig. 4E**). Decomposition of total energy into individual components revealed that engineered cores are characterized by improved van der Waals contacts, enhanced side-chain hydrogen bonding, and increased hydrophobic contributions consistent with the importance of tight side-chain packing within the fibril core. These gains are accompanied by penalties associated with burial of polar groups within the cores. Correlation analysis further supported this, revealing strong coupling between packing interactions and overall stability across the core rather than focused on specific residues, alongside trade-offs between polar and non-polar contributions (**Fig. 4F**). Consistent with these observations, analysis of sidechain and main-chain accessibility revealed modest but systematic improvement across designs (**Fig. 4G**), indicating that sequence variation modulates local packing density and solvent exposure without disrupting the overall architecture of the fibril. These findings support a non-unique mapping between sequence and structure in αS amyloid fibrils, where distinct energetic solutions can satisfy the same geometric constraints of the fold.

### *De novo* designs assemble into amyloid fibrils

To experimentally test the accuracy of the inverse folding approach for amyloid cores, we recombinantly expressed the selected *de novo* designed sequences and assessed their ability to assemble into amyloid fibrils. Of the nine selected designs, eight were successfully expressed and purified, while one construct (M18) could not be obtained in soluble form and was excluded from further analysis. Fibril formation was monitored under the same conditions used to determine the template fibril structure^17^. Transmission electron microscopy (TEM) revealed that all expressed constructs assembled into fibrillar aggregates characteristic of amyloid structures (**Fig. 5A**). Morphological analysis indicated differences among the self-assembled design fibrils. Three designs (M5, M6, and M19) self-assembled into fibrils with widths resembling those of WT αS under matching assembly conditions, whereas the remaining constructs formed morphologically distinct assemblies (**Fig. 5B**). Aggregation kinetics were monitored using thioflavin T (ThT) fluorescence assays. All designs exhibited sigmoidal aggregation kinetics characteristic of nucleation-dependent amyloid formation (**Fig. 5C**), but substantial variation in aggregation efficiency was observed across constructs. The three designs sharing morphological properties with αS fibrils, M5, M6, and M19, in addition to M11, displayed higher endpoint ThT amplitude (**Fig. 5D**), consistent with enhanced fibril formation under these conditions. M6 and M19, in addition to M1 and M9 also exhibited shorter half-times (t_1/2_) (**Fig. 5E**), indicating accelerated assembly kinetics relative to αS. To assess the thermodynamic stability of the fibrils formed by each design, we performed sedimentation experiments to estimate the critical concentration of soluble monomer at equilibrium. These measurements revealed differences in fibril stability among the designs, with M1, M16, and M19 similar stability to αS, and M5 exhibiting improved stability (**Fig. 5F**). Together, these results demonstrate that all sequences sampled from the compatibility space readily assemble into amyloid fibrils *in vitro*, with a subset of designs reproducing morphological features of the template strain or improved kinetics and fibril stability, linking sequence–structure compatibility to functional and programmable amyloid assembly.

**Figure 5.**
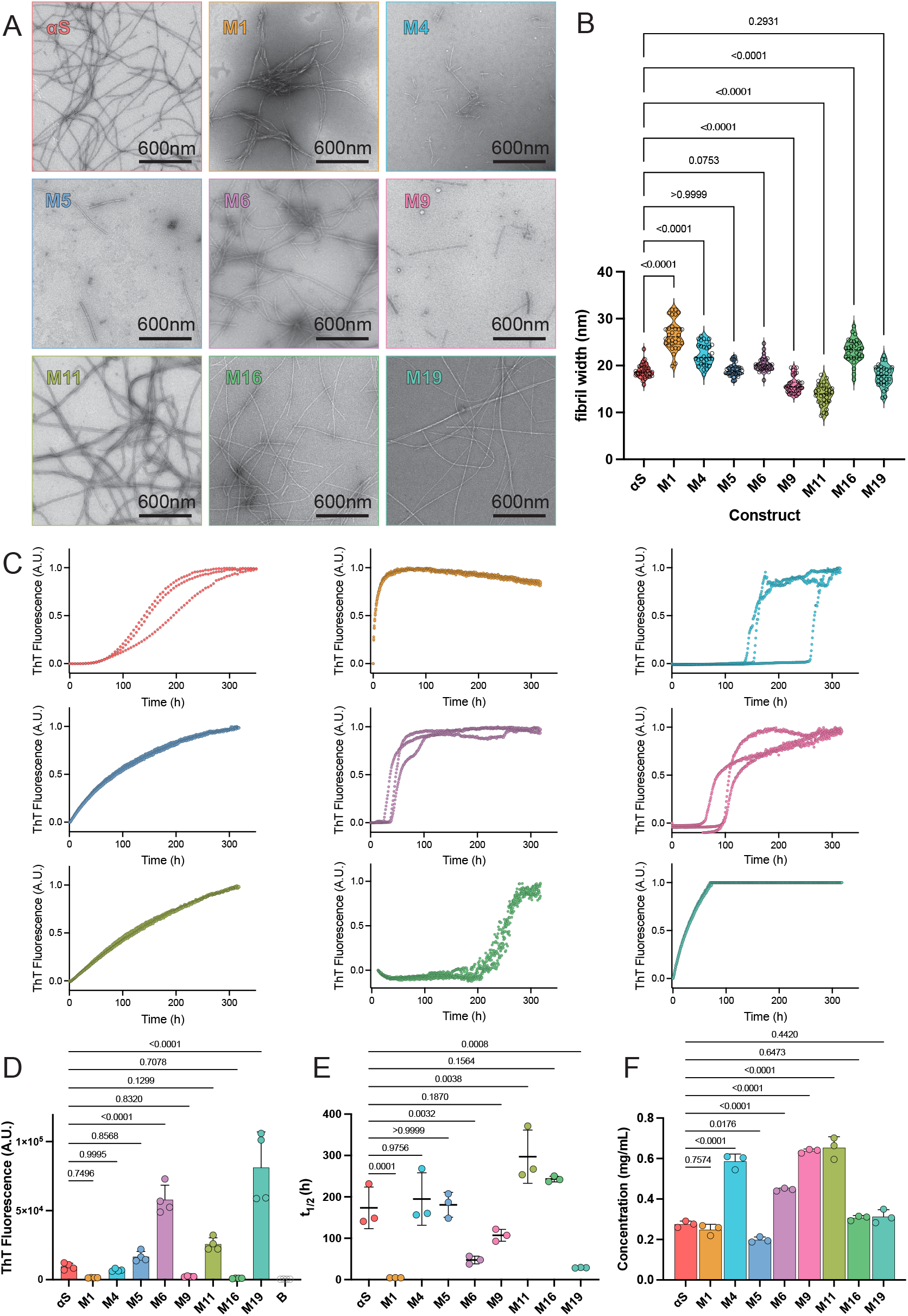
Characterization of amyloid fibrils formed by *de novo* designed sequences. (**A**) Electron micrographs of fibrils formed by αS and *de novo* designs in matching conditions. Scale bars, 600 nm. (**B**) Quantification of fibril widths across constructs (n=30 individual fibrils). Selected designs (M5, M6, M19) produce fibrils with widths comparable to αS. (**C**) Fluorescence aggregation kinetics for αS and *de novo* designs (n=3 independent repeats). End-point fluorescence intensities and (**E**) aggregation half-times derived from fluorescence assays. (**F**) Sedimentation analysis reflects differential stability of the designed constructs in equilibrium experiments (n=3 independent repeats). Statistics: One-way ANOVA with Dunnett correction for multiple comparisons.

### Spectral fingerprinting and cross-seeding reveal structural compatibility

To further assess whether fibrils formed by the designed sequences share structural features with αS fibrils, we performed spectral fingerprinting using the amyloid-sensitive dye pFTAA, which reports on fibril conformation through surface binding-dependent fluorescence emission profiles. Emission spectra recorded for each *de novo* design were compared to those of WT αS fibrils (**Fig. 6A**). Principal component analysis (PCA) revealed that fibrils formed by M5 and M19 cluster together with αS in the eigen space, indicating similar surface environments and underlying structural features (**Fig. 6B**). Fibrils formed by M6 also localized near this cluster, consistent with its morphological similarity to the WT assemblies, whereas the remaining designs occupied more distant regions of the embedding. To test this compatibility further, we next performed cross-seeding experiments using pre-formed fibrils from each design to initiate aggregation of αS monomer. A subset of designs (M5, M6, M11, and M19) efficiently seeded αS aggregation, significantly reducing the lag phase relative to unseeded reactions (**Fig. 6C**). Quantification of aggregation half-times confirmed strong cross-seeding activity for these constructs (**Fig. 6D**). Notably, these same designs correspond to those that exhibit spectral overlap with αS and produce fibrils with similar morphology, indicating convergence of independent readouts. In line with these observations, TEM revealed that cross-seeded fibrils adopt morphologies indistinguishable from self-seeded αS assemblies (**Fig. 6E**), with overlapping fibril width distributions (**Fig. 6F**). Together, these results demonstrate that a subset of the designed sequences produces fibrils that are structurally compatible with the WT αS strain and can template its assembly by leveraging cross-templating as a functional proxy.

**Figure 6.**
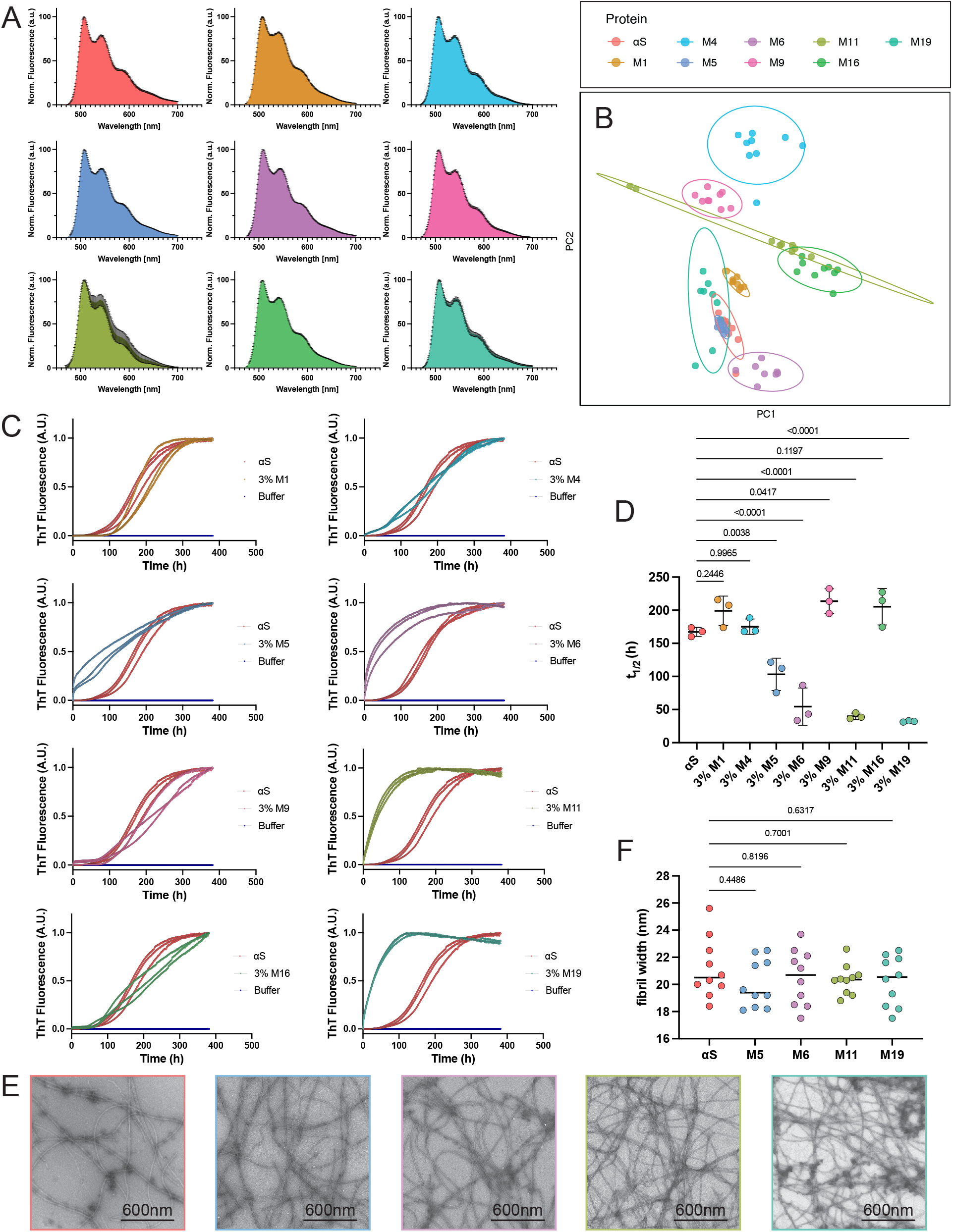
Structural compatibility of designed fibrils assessed by spectral fingerprinting and cross-templating. (**A**) pFTAA fluorescence emission spectra of fibrils formed by αS and *de novo* designs (n=9 independent repeats). (**B**) PCA of the collected emission spectra, showing clustering of spectra derived from selected designs (M5, M19) with spectra collected from αS fibrils. (**C**) Cross-seeding of αS using pre-formed fibrils from each design (n=3 independent repeats). (**D**) Quantification of aggregation half-times under cross-seeding conditions. (**E**) Electron micrographs of second-generation fibrils formed by cross-seeding monomeric αS protein. Scale bars, 600 nm. (**F**) Distribution of fibril widths for cross-seeded assemblies (n=10 individual fibrils). Statistics: One-way ANOVA with Dunnett correction for multiple comparisons.

### Functional validation of designs in cells

To determine whether structurally compatible designs retain the ability to propagate in a cellular context, we evaluated their seeding activity using αS (1–120) biosensor cells^22^. These biosensors provide a highly sensitive and selective readout of αS templated aggregation, with prior studies demonstrating that seeded aggregation in such systems depends strongly on the structural compatibility of the incoming fibril seed^21,22,44,45^. Pre-formed fibrils were introduced into cells, and seeded aggregation was monitored via FRET-based detection of inclusion formation (**Fig. 7A**). While most *de novo* designs failed to induce detectable aggregation, two constructs (M5 and M19) robustly seeded the WT αS biosensor cells (**Fig. 7B**). Quantification of aggregation levels showed that M19 induces aggregation comparable to αS fibrils (**Fig. 7C**), indicating efficient cellular propagation. Importantly, these constructs correspond to those that exhibit morphological similarity to αS fibrils and strong cross-templating activity *in vitro*, once again linking structural compatibility across independent assays. The inability of other designs to seed in cells, despite forming fibrils *in vitro*, potentially highlights the stringent requirements for productive templating in a cellular environment, consistent with observations both in αS as well as in other amyloid target biosensors^21,22,44,45^. Together, these results demonstrate that a subset of sequence-compatible designs gives rise to fibril assemblies that can propagate aggregation in a cellular environment, providing functional validation of the fibril reprogramming strategy.

**Figure 7.**
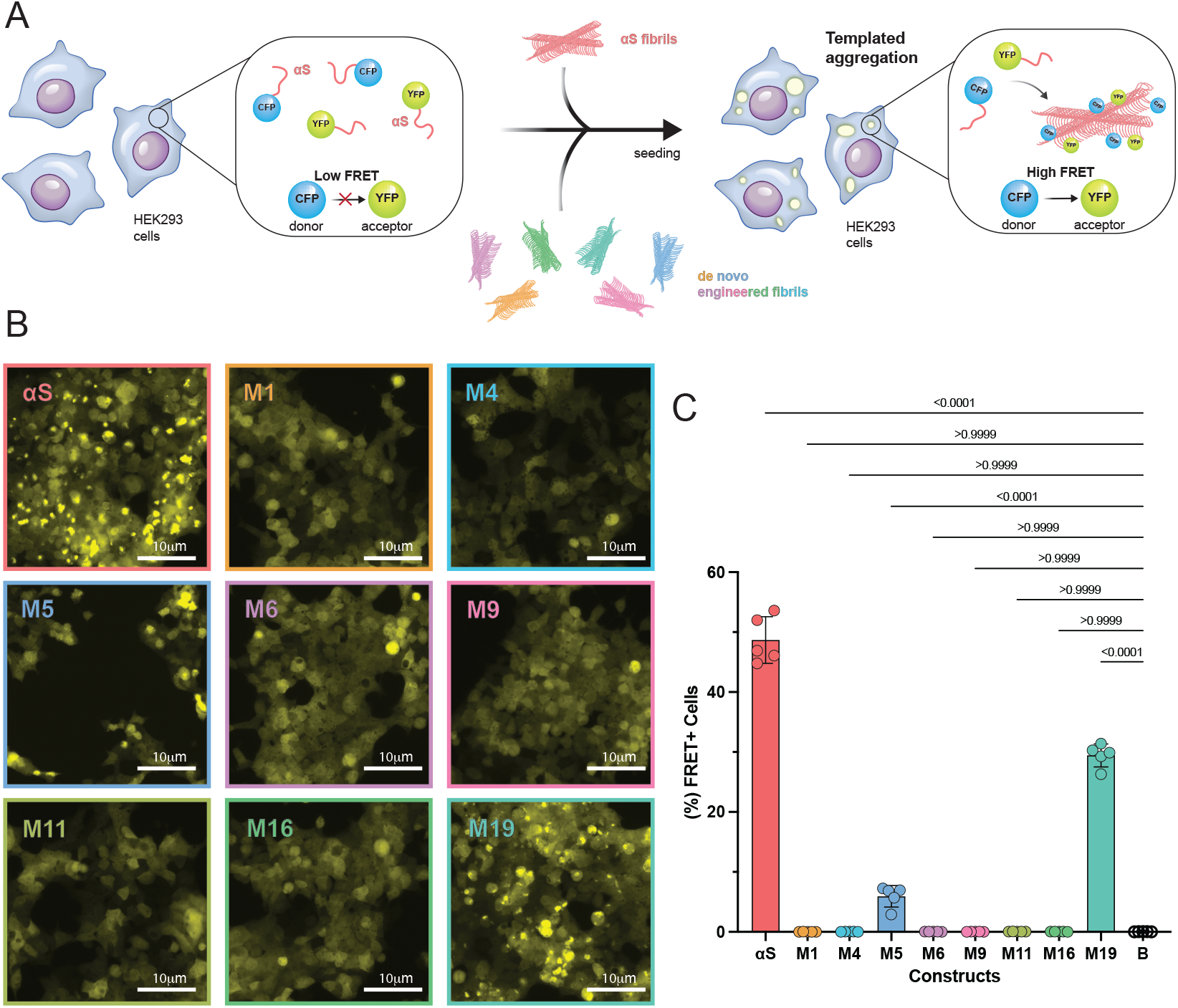
Functional validation of designed fibrils in αS biosensor cells. (**A**) Schematic of the αS biosensor assay for detecting seeded cross-templated aggregation. (B) Confocal imaging and (C) quantification of seeded aggregation in biosensor cells following treatment with fibrils derived from designs (n=5 independent repeats). M5 and M19 sequences exhibit robust seeding activity. Statistics: One-way ANOVA with Dunnett correction for multiple comparisons.

## Discussion

Amyloid fibrils display pronounced structural polymorphism, enabling a single protein sequence to give rise to distinct strains associated with several different proteinopathies. While considerable progress has been made in resolving the atomic structures of polymorphic amyloid fibril assemblies derived from several proteins^46^, the extent to which a given fibril fold constrains the underlying amino-acid sequence has remained unclear. While a single polypeptide sequence can give rise to multiple fibril polymorphs, our results demonstrate the converse: a given fibril architecture can be encoded by a diverse set of sequences. By combining structure-guided generative protein design with experimental validation, we show that αS amyloid folds define a continuous sequence compatibility landscape, within which multiple sequence solutions can give rise to structurally equivalent yet energetically distinct assemblies with convergent functional behaviour. Compatibility of sequences with a given αS fibril fold is not discrete but instead organized along a continuous manifold. Sampling of sequence space under increasing temperatures revealed gradual shifts in aggregation propensity and physicochemical properties, rather than abrupt transitions. This behaviour is reflected in the low-dimensional embedding of the designed sequences, which occupy a continuous region of sequence space with a clear directional drift. Importantly, although sequences sampled at higher temperatures become increasingly divergent from the αS sequence, they retain overall physicochemical characteristics compatible with the amyloid fold, indicating that substantial sequence variation can be accommodated without loss of structural compatibility.

At the energetic level, this compatibility arises from the ability of different sequences to satisfy the same structural constraints through distinct local interactions. Residue-level energetic decomposition revealed that stabilizing and destabilizing contributions are distributed throughout the fibril core, rather than localized to specific positions. Despite this heterogeneity, all designs maintain comparable overall stability within previously reported energy barriers compatible with αS cross-elongation^47^, indicating that multiple energetic solutions can converge to support a common structural framework. In particular, van der Waals interactions emerge as a dominant stabilizing force, consistent with the importance of steric zipper-like tight side-chain packing in the fibril core^48^, while electrostatic and solvation contributions exhibit compensatory behaviour. Together, these results support a non-unique mapping between sequence and structure in amyloid fibrils, in which diverse sequences can stabilize the same fold through different combinations of local interactions. These findings are consistent with an emerging view of amyloid aggregation as a degenerate energy landscape, in which multiple, potentially predictable^49^, sequence solutions can stabilize structurally related fibril architectures. Recent systematic mutational studies in αS fibrils have shown that perturbations within fibril core regions can reshape the energy landscape by introducing alternative aggregation pathways and structural minima, whereas mutations outside the core modulate kinetic barriers within a conserved framework^50,51^. In this context, our results extend these observations by demonstrating that such alternative sequence solutions can be accessed not only through targeted mutagenesis, but potentially through structure-guided generative design of a much wider sequence space. This highlights the ability of amyloid folds to accommodate diverse sequences within a constrained structural framework, while maintaining access to multiple energetically viable states.

Experimental characterization confirms that sequences sampled from this compatibility landscape can assemble into amyloid fibrils, while exhibiting diverse kinetic and structural properties. Although all tested *de novo* designs form fibrillar assemblies, their aggregation rates, fibril morphology, and stability vary substantially, indicating that sequence variation modulates the kinetic and thermodynamic features of fibril formation without abolishing the ability to assemble. These differences likely reflect variations in nucleation efficiency, elongation dynamics, and local packing interactions, which collectively shape the observed aggregation behaviour. Comparison of their physicochemical properties reveals that designs exhibiting faster kinetics or higher aggregate fractions tend to occupy a regime characterized by moderate aggregation propensity profiles and low to near-neutral hydropathy values, suggesting that efficient assembly does not arise from maximization of hydrophobicity but rather from a balanced interplay between solubility and intermolecular interactions. Spectroscopic analysis and cross-seeding experiments demonstrate that certain designs can form fibrils that are compatible with cross-templating WT aggregation and reproduce structural features characteristic of the native αS fibrils. These sequences exhibit shifts in secondary structure propensities, higher hydrophilicity, and higher disorder content implicated in modulating aggregation^52–54^, with minimal distortion across other physicochemical descriptors. In contrast, designs that aggregate robustly yet fail to cross-seed αS display pronounced deviations along individual physicochemical axes, including elevated amyloidogenic potential or shifts in secondary structure features. Such imbalances may introduce alternative aggregation hotspots, thereby redirecting assembly pathways and generating barriers to cross-templating. Together, the above suggest that while the amyloid fold tolerates substantial sequence variation, compatibility with a specific strain is constrained by coordinated interactions across multiple features, and that overrepresentation of individual descriptors can shift sequences outside the functional compatibility regime.

These observations are consistent with the concept of amyloid strains as structurally distinct yet partially overlapping states within a shared energy landscape^1,3,4^ and provide a direct connection to naturally occurring sequence perturbations in αS, including hereditary mutations and post-translational modifications. Familial mutations such as E46K, H50Q, G51D, and A53T have been shown to induce modest changes in fibril folds, with the overall architecture largely preserved^55–58^. This structural conservation is consistent with their ability to cross-seed αS, albeit often with reduced efficiency, likely reflecting subtle mismatches in local packing or interfacial interactions. In contrast, certain post-translational modifications can induce more pronounced structural changes. For example, O-GlcNAcylation promotes fibril polymorphs with altered topology and reduced seeding capacity^59^, and phosphorylation at Y39 stabilizes distinct conformational states with altered aggregation behaviour, while acetylation of lysine residues modulates aggregation and seeding capacity of αS^60–63^. In line with the above, our findings support a model in which local sequence perturbations can either preserve the overall energy landscape, modulating barriers within a WT-like framework and extending prior work with short fragments that can recapitulate key structural features of fibril cores^64–66^, or introduce new minima corresponding to alternative fibril polymorphs. Finally, assessment of seeding activity in cellular biosensor assays reveals that while some designs that are compatible with the WT structure *in vitro* retain seeding activity in cells, others do not, suggesting that additional factors, such as fibril stability, surface properties, or interactions with cellular components, may contribute to propagation efficiency^67–70^.

Beyond implications for disease mechanisms, our findings have broader relevance for the emerging field of amyloid engineering. Previous studies have demonstrated that aggregation-prone regions can be rationally redesigned to modulate protein aggregation, including engineering enhancers^71,72^ or inhibitors of protein aggregation^73–77^. In parallel, the structural robustness of amyloid assemblies has been exploited to engineer highly stable biomaterials, including β-solenoid scaffolds^78,79^, cross-linked fibrillar hydrogels^80^, and functional nanomaterials such as conductive fibers^81^, filtration systems^82^, and catalytic assemblies^83–86^. These approaches have largely relied on local sequence modifications of existing scaffolds. Extending this concept further, our results demonstrate that structure-guided generative design enables exploration of the broader combinatorial sequence space compatible with a defined amyloid fold. This provides a framework for designing amyloid assemblies with tailored structural and functional properties, extending beyond incremental mutation-based strategies toward a more global understanding and programmable control over amyloid sequence–structure relationships.

In summary, our results establish that αS fibrils are governed by a constrained but continuous sequence compatibility landscape, in which diverse sequences can encode a common structural framework while giving rise to distinct functional outcomes. By integrating generative design, energetic analysis, and experimental validation, we provide a framework for systematically probing and engineering amyloid sequence–structure relationships. This approach opens new avenues for understanding the molecular determinants of amyloid polymorphism and strain behaviour, and for harnessing amyloid assemblies as programmable biological and materials systems.

## Materials and Methods

### Structural Templates and Model Preparation

Cryo-EM structures of αS fibrils representing distinct structural polymorphs were used as design templates, including an MSA-derived model (PDB ID: 6XYO), an *in vitro* fibril structure (PDB ID: 6H6B), and a patient-derived fibril structure (PDB ID: 8A9L) (**Fig. 2A**). For each template, fibril coordinates were processed to generate a multi-layer assembly representing a minimal repeating unit along the fibril axis. To enable consistent sequence design and energetic evaluation, structures were normalized to a six-chain representation, in which chains were aligned, ordered along the fibril axis, and assigned consistent identifiers. This ensured structural equivalence across layers and enabled symmetry-constrained sequence design.

### Deep-learning Sequence Design

Sequence design was performed using inverse folding with ProteinMPNN^28^ (**Fig. 1**). For each template, all chains within the fibril assembly were designed simultaneously under a symmetry constraint, such that identical amino acid identities were enforced across equivalent stacked positions in each chain. Sequence design was restricted to the A segment of the fibril core, while solvent-exposed positions (to preserve surface properties) and the B segment were held fixed. The resulting design region corresponds to a structurally ordered but less densely packed portion of the fibril core, providing a larger accessible sequence space under the constraints of the target fold. In contrast, residues within the B segment, which overlap extensively with the NAC region, were excluded from design. Additionally, the A segment exhibits higher geometric conservation across distinct αS fibril structures, enabling more consistent design constraints across templates. Sequence sampling was performed across a temperature grid (T = 0.1, 0.2, 0.3, 0.4, 0.5, 0.8, 1.0, and 1.2), which modulates the degree of sequence exploration during sampling. For each temperature, 500 sequences were generated per template, resulting in a total of 4000 sequences per structure. Logos of multiple sequence alignments of the derived designs were generated using Logomaker (v0.8.7)^87^.

### Energetic Analysis and Thermodynamic Profiling

Designed sequences were subjected to energetic evaluation using PyRosetta^88^. Each sequence was threaded onto the corresponding fibril template and subjected to structural relaxation using Cartesian FastRelax with the ref2015_cart energy function. Relaxed structures were scored, and sequences were ranked based on total Rosetta energy. Top-ranking candidates were selected for further structural refinement and downstream analysis.

Energetic analysis was performed using the FoldX force field^89^. All models were generated using the αS fibril template (PDB ID: 6H6B). Specifically, the top-ranking designs were introduced into the αS fibril structure using the BuildModel function. This approach generates mutant structures by substituting amino acids while preserving the backbone conformation of the template and optimizing local side-chain packing. Residue-level energetic contributions were calculated using the SequenceDetail function in FoldX. This analysis provides per-residue decomposition, including van der Waals interactions, backbone and side-chain hydrogen bonding, electrostatics, solvation (polar and hydrophobic), steric clashes, and burial values. Global energetic comparisons (ΔΔG values) across designs were quantified by subtracting aggregated template energetics from the individual energy components of each design. To assess relationships between different energetic contributions, residue-level individual energy components were analysed using pairwise correlation plots. Correlation matrices were generated in R using the ggpairs function.

Finally, thermodynamic profiling (per-residue fibril stability contributions) was quantified using our previously established thermodynamic profiling pipeline^2,4^, in which all inter-residue interaction energies within the amyloid cores were computed and summed to yield per-residue stabilization contributions. Structural inspection, model manipulation, and molecular graphics were performed in ChimeraX (v1.10.1) and Pymol (v3.1.8). All statistical analyses and data visualization were carried out in GraphPad Prism (v10.6.1).

### Sequence Propensity Analysis

Aggregation propensity of all designed sequences was evaluated using TANGO^41^ and WALTZ^42^ predictors. TANGO scores were used to estimate general β-sheet aggregation propensity driven by hydrophobic interactions, while WALTZ scores were used to identify sequence-specific amyloid-forming motifs. Both total scores and aggregation-prone windows were computed for each sequence. Secondary structure changes were calculated using the Chou-Fasman propensities^90^ and hydropathy analysis was performed using Protein GRAVY, which returns the GRAVY (grand average of hydropathy) value for protein sequences. The GRAVY value is calculated by adding the hydropathy value for each residue^91^ and dividing by the length of the sequence. Sequence-level physicochemical descriptors were calculated for each sequence, including amino acid composition, blosum62 distances to the WT sequence, and residue class fractions in R. To characterize the structure of the sampled sequence space, sequences were embedded into a reduced-dimensional representation using Uniform Manifold Approximation and Projection (UMAP) based on physicochemical and aggregation-related features, using the uwot package in R.

### Protein Expression and Purification

Expression and purification of αS (residues 1-121) and *de novo* designs (M1, M4, M5, M6, M9, M11, M16, and M19) were performed following the protocol described previously^17^, with minor modifications. Gene fragments encoding the *de novo* engineered designs and WT αS (1–121) were synthesized (Twist Bioscience), cloned into the pET28a expression vector, and transformed into Escherichia coli BL21(DE3) cells for protein expression. Cultures were grown in LB medium supplemented with kanamycin (50 mg/L), at 37 °C with shaking at 200 rpm to an OD600 of 0.6– 0.9. Protein expression was induced by the addition of 1 mM IPTG, followed by incubation for 4 h at 37 °C under the same shaking conditions. Cells were harvested by centrifugation at 5000 rpm for 10 min and resuspended in lysis buffer containing 50 mM Tris-HCl, 150 mM NaCl, and 10 mM EDTA (pH 8.0). A protease inhibitor cocktail (Roche) was added to prevent proteolytic degradation during purification. Cells were lysed on ice using an Omni Sonic Ruptor 400, and the lysate was clarified by centrifugation to remove cellular debris. The clarified supernatant was applied to HisPur™ Ni-NTA resin (Thermo Fisher Scientific) for affinity purification. The column was washed with 20 mM sodium phosphate buffer (pH 7.4) to remove non-specifically bound proteins, and the target protein was eluted with 250 mM imidazole. Eluted protein fractions were concentrated and buffer-exchanged into Dulbecco’s phosphate-buffered saline (DPBS; Gibco: 2.66 mM KCl, 1.47 mM KH_2_PO_4_, 137.93 mM NaCl, and 8.06 mM Na_2_HPO_4_, pH 7.3) using a spin column. Purified proteins were stored at −20 °C until further use. Protein purity was assessed by SDS–PAGE analysis.

### SDS-PAGE

Protein samples (10 µL) were mixed with 4x Laemmli Buffer (Bio-Rad) containing 2-mercaptoethanol and heated at 95°C for 5 min. The samples were then loaded and resolved on NuPAGE™ 4-12% Bis-Tris gels (Thermo Fisher). Precision Plus Protein™ molecular weight ladder (Bio-Rad) was used as a size marker. The gels were stained with SimplyBlue™ safestain for ~1h and destained in distilled water. Gel images were acquired using the Syngene G:BOX imaging system.

### Preparation of Amyloid Fibrils

Amyloid fibrils were generated from recombinant protein at 5mg/mL following a previously published protocol^17^. Protein solutions (200 µL) were incubated (DPBS, Gibco; 2.66 mM KCl, 1.47 mM KH_2_PO_4_, 137.93 mM NaCl, 8.06 mM Na_2_HPO_4_, pH 7.0–7.3) at 37°C with constant shaking at 1000 rpm in an orbital thermomixer (Eppendorf). Reactions were allowed to proceed for 5 days to ensure fibril formation.

### Thioflavin T (ThT) fluorescence assays

Aggregation kinetics were monitored using ThT fluorescence assays. Protein samples (2 mg/mL) were incubated in 96-well clear-bottom plates (Corning) at 37°C with continuous shaking at 300 rpm in the presence of 25 µM ThT. Fluorescence changes were measured at regular intervals using excitation and emission wavelengths at 448 nm and 482 nm, respectively, in a FLUOstar Omega (BMG LABTECH) plate reader. Data analysis was performed with BMG Labtech’s Mars Omega, and the obtained data were fitted using GraphPad Prism (V 11.0.0).

### Fluorescence Spectroscopy

Preformed fibrils from WT αS or *de novo* designs were diluted to 50 µM and incubated with 0.5 µM the oligothiophene dye pentameric formyl thiophene acetic acid (pFTAA). Emission spectra were recorded between 468 and 700 nm following excitation at 440 nm. Spectral data were analysed using principal component analysis (PCA) with the PCA function in R to compare conformational differences between fibril samples.

### Transmission Electron Microscopy (TEM)

Fibril samples were applied (5 µL) to glow-discharged carbon-coated grids (400-mesh; Electron Microscopy Sciences) and incubated for 5 min at room temperature. Grids were washed with Milli-Q water (5 µL) to remove excess samples and stained with 2% (w/v) uranyl acetate for 2 min. The excess stain was removed using filter paper. After air-drying, grids were imaged using a transmission electron microscope (JEOL1400) operating at 80kV. Fibrils widths were quantified from individual images (n=3) using ImageJ (v21.0.7).

### Solubility Assay

Following fibril formation, samples were diluted to 1mg/ml and centrifuged at 100,000 × g for 45 min at 4 °C to separate soluble and insoluble fractions. The concentration of soluble protein in the supernatant was determined by absorbance at 280 nm using a DeNovix DS-11 spectrophotometer.

### Cross-seeding Experiments

Preformed fibrils were diluted to 2 mg/mL and sonicated to generate seeds. Fibril suspensions were sonicated at 20% amplitude for 3 min (3 s ON, 1 s OFF) at 4 °C using a probe sonicator. Cross-seeding reactions were performed by adding 3% (v/v) preformed fibril seeds (WT or *de novo* designs) to monomeric WT αS (2 mg/mL) in DPBS containing 25 µM ThT. Seed-free WT samples were used as controls. Samples were incubated at 37 °C with shaking (300 rpm), and aggregation kinetics were monitored by ThT fluorescence using excitation and emission wavelengths at 448 nm and 482 nm, respectively, in a FLUOstar Omega (BMG LABTECH) plate reader.

### Cell Biosensor Assays

αS biosensor cells expressing WT αS (residues 1–120) fused to C-terminal CFP and YFP fluorescent tags^22^ were plated in 96-well plates (15000 cells per well) and cultured in 80 mL DMEM supplemented with 10% FBS, 1% Penicillin/Streptomycin and 1% Glutamax in a 96-well plate and incubated at 37°C. Fibril seeds were generated by sonication (65% amplitude, 30 s ON/30 s OFF cycles for 8 min) and delivered to cells using lipid-mediated transfection, mixed with Lipofectamine 2000 (0.5 µL per well) in Opti-MEM. After 48 h, cells were harvested (Accumax) and fixed (1% paraformaldehyde in PBS). The cells were pelleted and resuspended in PBS containing 3% FBS, then analysed by flow cytometry to quantify fluorescence resonance energy transfer (FRET) as a measure of intracellular aggregation, in an Attune NxT flow cytometer using excitation: 405nm, emission:450/50 nm (For CFP); excitation: 488nm emission: 530/30nm (For YFP), excitation: 405nm, and emission: 512/25nm for FRET. For identifying FRET-positive cells, HEK cells were first gated on FSC versus SSC to isolate the main cell population. From this population, single cells were selected using FSC-A versus FSC-H to exclude doublets and multiplets. Within the singlet gate, double-positive cells were identified based on CFP and YFP fluorescence. FRET-positive events were defined as double-positive cells exhibiting increased FRET signal relative to negative control. Data was processed in FlowJo (version 10.10.1).

## Acknowledgements

This work was supported by the National Institute on Aging (NIA) of the National Institutes of Health under award number [1RF1AG078888-01] to LAJ and by a scholarship from the Thomas O. Hicks Scholar in Medical Research awarded to NL. KK was partially supported by the O’Donnell Brain Institute (OBI) Sprouts grant program. Computational resources were provided by the BioHPC cluster supported by the Lyda Hill Department of Bioinformatics at UTSW. We acknowledge the UT Southwestern Electron Microscopy Core, funded by the NIH grants [1S10OD021685-01A1].

## Author contributions

LG and KK performed the experiments. AM, JC and NNL performed computational analysis. LJ and NNL conceived and initiated the study. LG, KK, JC, and NNL performed data analysis. LJ, and NNL provided resources and funding. NNL supervised the project. LG and NNL prepared the original draft, and all authors contributed to the review and editing of the final version of this manuscript.

## Data Availability statement

All experimental and computational raw data generated are available from the corresponding author on reasonable request.

